# Identification and functional characterization of two CXCL17 paralogs from zebrafish

**DOI:** 10.1101/2025.08.27.672743

**Authors:** Jie Yu, Wen-Feng Hu, Juan-Juan Wang, Ya-Li Liu, Zeng-Guang Xu, Zhan-Yun Guo

## Abstract

C-X-C motif chemokine ligand 17 (CXCL17) and its receptor G protein-coupled receptor 25 (GPR25) have been identified as a significant pair in regulating immunity, but CXCL17 orthologs have not yet been identified from non-mammalian vertebrates. This study aimed to identify and characterize non-mammalian CXCL17 orthologs based on key features of mammalian CXCL17s, such as a C-terminal Xaa-Pro-Yaa motif, a signal peptide, and six cysteine residues. Two possible CXCL17 paralogs were identified from zebrafish (*Danio rerio*): Dr-CXCL17 (encoded by *zgc:158701*) and Dr-CXCL17-like (encoded by *si:dkey-112a7.5*). Both are previously uncharacterized proteins with unknown functions because they lack overall sequence similarity to known mammalian CXCL17s. For functional characterization, recombinant Dr-CXCL17 and Dr-CXCL17-like were prepared via overexpression in *Escherichia coli* and subsequent *in vitro* refolding and their activity was tested using NanoLuc Binary Technology (NanoBiT)-based β-arrestin recruitment assays, NanoBiT-based ligand□receptor binding assays, and chemotaxis assays. The results showed that both Dr-CXCL17 and Dr-CXCL17-like tightly bound to and efficiently activated zebrafish GPR25 (Dr-GPR25) and induced chemotactic movement in transfected human embryonic kidney (HEK) 293T cells expressing the receptor. Deletion of three C-terminal residues in both paralogs almost eliminated their binding, activation, and chemotactic effects, which suggests that this fragment is crucial for their function. Homologs of Dr-CXCL17 or Dr-CXCL17-like were retrieved from several other ray-finned fish species, indicating that two CXCL17 paralogs are present in certain fish species and function as endogenous agonists for the fish GPR25 receptor. The identification of fish CXCL17 orthologs suggests that the CXCL17□GPR25 pair likely originated in ancient fishes and was conserved across vertebrate lineages. This work represents the first identification of CXCL17 orthologs in non-mammalian vertebrates, paving the way for future functional studies of this ligand□receptor pair.

## Introduction

C-X-C motif chemokine ligand 17 (CXCL17) functions as a chemoattractant for certain leukocytes, such as T cells, monocytes, macrophages, and dendritic cells [1□14]. It is also implicated in tumor development, likely through its regulation of tumor immunity [15□21]. However, the identity of its receptor remains controversial. The orphan G protein-coupled receptor 35 (GPR35) was first reported as its receptor in 2015 [22], but later studies did not support this pairing [23□25]. In recent years, both the chemokine receptor CXCR4 and the orphan MAS-related receptor MRGPRX2 were reported as its receptors [25,26]. Most recently, Ocón’s group and our group independently identified the orphan G protein-coupled receptor 25 (GPR25) as its receptor [27,28]. As a rarely studied A-class G protein-coupled receptor (GPCR), GPR25 is primarily expressed in specific immune cells, such as T cells, plasma cells, and B-cells, according to data from the Human Protein Atlas (https://www.proteinatlas.org). The fact that these GPR25-expressing leukocytes can be attracted by mucosal tissue-expressed CXCL17 [27] suggests that the CXCL17–GPR25 pair plays an important role in immune regulation.

According to the National Center for Biotechnology Information (NCBI) gene database, GPR25 orthologs are broadly distributed from fishes to mammals and exhibit substantial amino acid sequence similarity, suggesting that GPR25 serves important functions across all vertebrates. In contrast, CXCL17 orthologs have not been identified in non-mammalian vertebrates, implying that CXCL17 may be restricted to mammals. Consequently, the endogenous agonists of non-mammalian GPR25s remain unknown. A previous study reported that apelin and apela, two peptide hormones highly conserved among vertebrates, act as weak agonists for certain non-mammalian GPR25s [29]. Our recent work demonstrated that human CXCL17 exhibits markedly higher activity than apelin or apela toward fish GPR25s from both the ray-finned zebrafish (*Danio rerio*) and the lobe-finned coelacanth (*Latimeria chalumnae*) [30]. Based on these findings, we hypothesized that CXCL17 orthologs may indeed exist in non-mammalian vertebrates but remain unrecognized due to the absence of overall sequence similarity with known mammalian CXCL17s.

In this study, we identified and functionally characterized two zebrafish CXCL17 paralogs, designated as Dr-CXCL17 and Dr-CXCL17-like, based on key sequence features shared with mammalian CXCL17s. Both proteins were previously uncharacterized and of unknown function, as their lack of overall amino acid sequence similarity with mammalian CXCL17s precluded their recognition as CXCL17 orthologs. Recombinant Dr-CXCL17 and Dr-CXCL17-like were shown to bind to and activate zebrafish GPR25 (Dr-GPR25) and to induce chemotactic migration of transfected human embryonic kidney (HEK) 293T cells via their conserved C-terminal fragment, indicating that they act as endogenous agonists of GPR25 in zebrafish. Database searches further identified fish homologs through BLAST analysis using Dr-CXCL17 or Dr-CXCL17-like, suggesting that these two paralogs are present in certain fish species. To the best of our knowledge, this is the first report of CXCL17 orthologs in non-mammalian vertebrates, providing a foundation for future functional studies of the CXCL17–GPR25 pair in these species.

## Results

### Identification of possible fish CXCL17s

Our recent studies have shown that three C-terminal residues are essential for human CXCL17 to activate human GPR25, as well as zebrafish and coelacanth GPR25s [28,30]. These residues are highly conserved among mammalian CXCL17s, particularly the penultimate proline (Pro) residue (Fig. S1 and Table S1). We therefore hypothesized that, if present, fish CXCL17 orthologs would possess three analogous residues at their C-terminus. Based on this, we proposed a potential C-terminal motif for fish CXCL17s: Xaa-Pro-Yaa, where Xaa and Yaa are typically large aliphatic residues such as leucine (Leu), isoleucine (Ile), methionine (Met), or valine (Val). This motif served as a primary criterion for searching publicly available databases for candidate fish CXCL17s. In addition, we applied two supplementary criteria: (1) they should be secretory proteins, possessing an N-terminal signal peptide; and (2) their mature peptides should contain six cysteine (Cys) residues and be fewer than 200 amino acids in length.

Zebrafish (*Danio rerio*) is a widely used model organism whose genome has been fully sequenced, extensively analyzed, and well annotated. We retrieved all zebrafish secretory proteins with fewer than 200 amino acids from the UniProt database (https://www.uniprot.org/uniprotkb?query=Danio+rerio&facets=model_organism%3A7955%2Cproteins_with%3A49%2Clength%3A%5B1+TO+200%5D), and searched for potential CXCL17s among them using the proposed criteria. Among the 795 proteins obtained, two candidates were identified with UniProt IDs B0S594 and A0AB13AB19, respectively.

The genetic information for both zebrafish proteins is available in the NCBI gene database (Table S2 and S3). B0S594 is encoded by the gene *zgc:158701* (Gene ID: 100151367), located between *pafah1b3* and *ceacam1* on zebrafish chromosome 16 (Fig. S2). This gene produces a single mRNA (NM_001144821) and encodes a small secretory protein (NP_001138293) comprising 109 amino acids. A0AB13AB19 is encoded by the gene *si:dkey-112a7.5* (Gene ID: 100536854), situated between *ponzr5* and *ugt5g1* on zebrafish chromosome 7 (Fig. S3). This gene yields two mRNA transcripts (NM_001386806 and XM_073906074), both of which encode the same protein (NP_001373735 or XP_073762175) containing 95 amino acids. For clarity, in this study we designated B0S594 as Dr-CXCL17 and A0AB13AB19 as Dr-CXCL17-like.

As shown in Fig. 1A, neither Dr-CXCL17 nor Dr-CXCL17-like exhibits significant overall sequence similarity to human CXCL17 (Hs-CXCL17) or other known mammalian CXCL17s, and thus cannot be identified through sequence BLAST searches using mammalian CXCL17s as queries. Consequently, both proteins were previously uncharacterized and of unknown function. In contrast to the high sequence variability observed among CXCL17s, zebrafish GPR25 (Dr-GPR25) shares considerable sequence similarity with human GPR25 (Hs-GPR25) (Fig. 1B), despite the evolutionary divergence of humans and fishes approximately 400 million years ago.

**Fig. 1.**
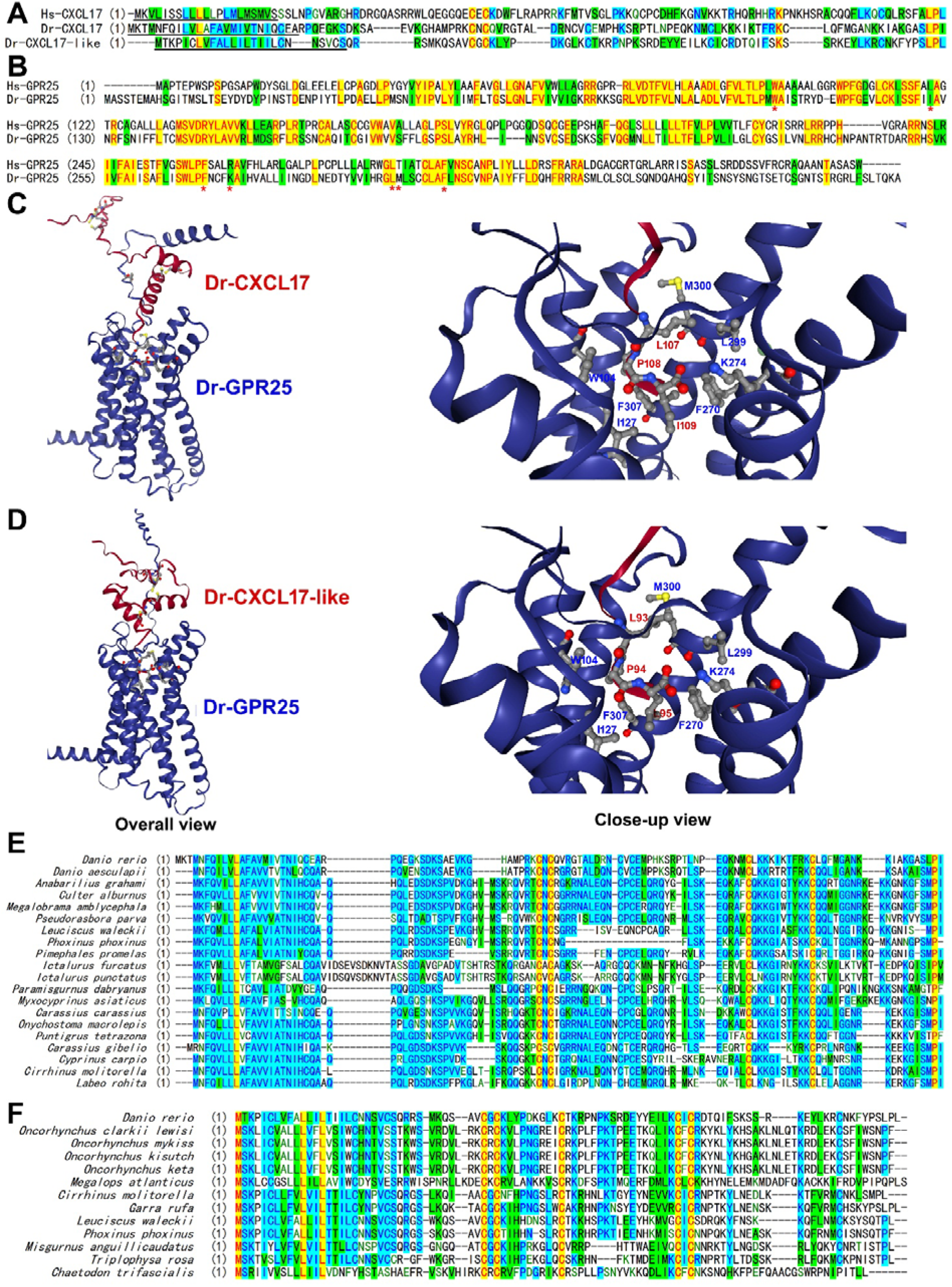
Identification of possible fish CXCL17s. (**A**) Amino acid sequence alignment of CXCL17s from human or zebrafish. These proteins were retrieved from NCBI database (Table S1□S3) and aligned via AlignX algorithm using the Vector NTI 11.5.1 software. Their signal peptide was predicted by SignalP-6.0 algorithm (https://services.healthtech.dtu.dk/services/SignalP-6.0/) and underlined. (**B**) Amino acid sequence alignment of GPR25s from human or zebrafish. Hs-GPR25 (NP_005289) and Dr-GPR25 (XP_073772858) were aligned via AlignX algorithm using the Vector NTI 11.5.1 software. The Dr-GPR25 residues involving ligand-binding are indicated by red asterisks. (**C,D**) AlphaFold3-predicted binding of the mature Dr-CXCL17 (C) and Dr-CXCL17-like (D) with Dr-GPR25. The binding structures were predicted via the online AlphaFold3 server (https://alphafoldserver.com) and viewed by an online server (https://nglviewer.org/ngl). The six Cys residues and three C-terminal residues of Dr-CXCL17 and Dr-CXCL17-like are shown as balls-and-sticks. The residues of Dr-GPR25 involving ligand-binding are shown as sticks-and-balls and indicated in panel B by red asterisks. In the predicted possible interactions are summarized in Table S4. (**E,F**) Amino acid sequence alignment of the fish homologs of Dr-CXCL17 (E) or Dr-CXCL17-like (F). These homologs were retrieved from the NCBI database via blast with Dr-CXCL17 or Dr-CXCL17-like (https://blast.ncbi.nlm.nih.gov), and their information is shown in Table S2 and S3. Some fish CXCL17 homologs have two variants (Table S2) due to alternative splicing and only the longer ones were aligned in panel D. The sequence alignment was conducted via AlignX algorithm using the Vector NTI 11.5.1 software.

The full-length Dr-CXCL17 contains a predicted N-terminal signal peptide of 25 residues and a mature peptide of 84 amino acids, whereas the full-length Dr-CXCL17-like has a predicted N-terminal signal peptide of 25 residues and a mature peptide of 70 amino acids (Fig. 1A). Both proteins contain six cysteine (Cys) residues in the mature peptide; however, their Cys arrangements differ: CXC-CXC-C-C in Dr-CXCL17 and CXC-C-CXC-C in Dr-CXCL17-like (Fig. 1A). The Cys pattern of Dr-CXCL17 is identical to that of mammalian CXCL17s, while the arrangement in Dr-CXCL17-like is distinct. The mature forms of both proteins are basic, with predicted isoelectric point (pI) values of 10.5 and 10.0, respectively. This basic nature may underlie their chemoattractant activity, as it could facilitate binding to negatively charged cell surface glycosaminoglycans after secretion, thereby forming local concentration gradients to attract immune cells.

According to AlphaFold2 predictions (AF-B0S594-F1-model_v4), Dr-CXCL17 adopts a flexible structure stabilized by three disulfide bonds (C47–C62, C49–C60, and C80–C91), but the overall prediction confidence is low. AlphaFold3 modeling further predicts that Dr-CXCL17 binds to Dr-GPR25 with an ipTM value of approximately 0.6, with the ligand’s C-terminal fragment inserted into the orthosteric ligand-binding pocket of the receptor (Fig. 1C). The three conserved C-terminal residues (I109, P108, and L107) of the ligand form extensive interactions with counterpart residues of Dr-GPR25 (Fig. 1C and Table S4). For example, the negatively charged C-terminal carboxyl moiety of Dr-CXCL17 forms electrostatic interactions with the positively charged ε-amine moiety of the receptor’s K274 residue; the penultimate P108 of the ligand forms hydrophobic interactions with the aromatic W104 of the receptor (Fig. 1C and Table S4).

AlphaFold3 predictions indicate that the mature Dr-CXCL17-like (residues 26–95) adopts a compact globular structure stabilized by three disulfide linkages (C36–C66, C38–C48, and C68–C86) (Fig. 1D). AlphaFold3 modeling also predicts binding of Dr-CXCL17-like to Dr-GPR25 with an ipTM value of approximately 0.6, in which the ligand’s C-terminal fragment inserts into the orthosteric ligand-binding pocket of the receptor (Fig. 1D). The three conserved C-terminal residues (L95, P94, and L93) of Dr-CXCL17-like form extensive interactions with counterpart residues of the receptor (Fig. 1D and Table S4). According to the AlphaFold3 predictions, Dr-CXCL17 and Dr-CXCL17-like have similar interaction patterns with receptor Dr-GPR25 (Table S4).

Amino acid sequence BLAST searches using Dr-CXCL17 identified several fish homologs in publicly available databases (Fig. 1E and Table S2). These homologs typically possess an N-terminal signal peptide for secretion, a mature peptide containing six cysteine residues arranged in the CXC-CXC-C-C pattern, and three highly conserved C-terminal residues. Similarly, BLAST searches using Dr-CXCL17-like retrieved homologs from certain fish species (Fig. 1F and Table S3), which feature an N-terminal signal peptide and a mature peptide with six cysteine residues arranged in the CXC-C-CXC-C pattern. All of these fish CXCL17 and CXCL17-like homologs remain uncharacterized and their functions are unknown.

### Preparation of the zebrafish CXCL17s via bacterial overexpression

To rapidly produce Dr-CXCL17 and Dr-CXCL17-like for functional characterization, we employed a bacterial overexpression strategy previously used for human CXCL17 (Hs-CXCL17) in our recent studies [28,30]. For purification purposes, a 6×His tag was fused to the N-terminus of the mature peptide of each zebrafish protein (Fig. S4). Sodium dodecyl sulfate-polyacrylamide gel electrophoresis (SDS-PAGE) analysis revealed that both 6×His-Dr-CXCL17 and 6×His-Dr-CXCL17-like were expressed in *Escherichia coli* predominantly as inclusion bodies, with particularly high accumulation observed for 6×His-Dr-CXCL17-like (Fig. 2A,B). Following solubilization of the inclusion bodies using an *S*-sulfonation approach, the proteins were purified via immobilized metal ion affinity chromatography (Ni²□ column). SDS-PAGE analysis of the eluted fractions showed the expected monomeric form (indicated by an asterisk) along with larger oligomers (Fig. 2A,B), indicating a tendency of the zebrafish CXCL17s to undergo intermolecular cross-linking, a phenomenon also observed for recombinant Hs-CXCL17 [28].

**Fig. 2.**
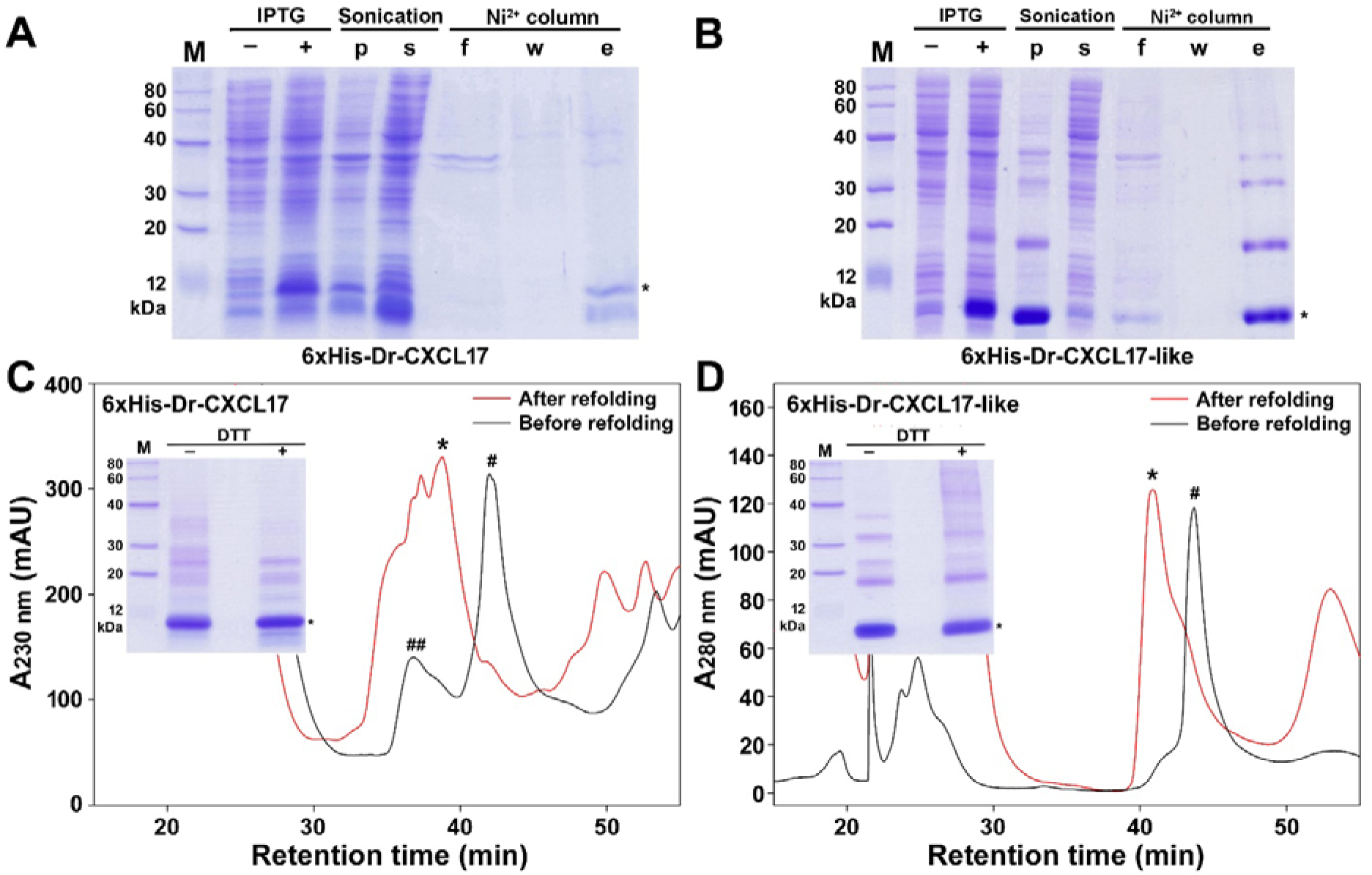
Preparation of the zebrafish CXCL17 and CXCL17-like via bacterial overexpression. (**A,B**) SDS-PAGE analysis of the samples of 6×His-Dr-CXCL17 (A) and 6×His-Dr-CXCL17-like (B). Lane (M), protein ladder; lane (□), before IPTG induction; lane (+), after IPTG induction; lane (p), pellet after sonification; lane (s), supernatant after sonification; lane (f), flowthrough from the Ni^2+^ column; lane (w), washing fraction by 30 mM imidazole; lane (e), eluted fraction by 250 mM imidazole. After electrophoresis, the SDS-gel was stained by Coomassie brilliant blue R250. Band of the monomeric zebrafish CXCL17 ortholog was indicated by an asterisk. (**C,D**) HPLC analysis of the samples of 6×His-Dr-CXCL17 (C) and 6×His-Dr-CXCL17-like (D). The major peak (indicated by an asterisk) after refolding was analyzed by SDS-PAGE and used for activity assays. **Inner panel**, SDS-PAGE analysis of the major refolding peak. Lane (M), protein ladder; lane (□), without DTT treatment; lane (+), with DTT treatment. After electrophoresis, the SDS-gel was stained by Coomassie brilliant blue R250.

The eluted fractions from the Ni²□ column were further analyzed by high performance liquid chromatography (HPLC). For both proteins, a broad major peak (indicated by a hash symbol) was eluted from a C_8_ reverse-phase column (Fig. 2C,D, black trace). In the case of 6×His-Dr-CXCL17, a broad minor peak (indicated by a double hash symbol) was also detected (Fig. 2C), likely representing partially degraded products, as suggested by SDS-PAGE analysis (Fig. 2A). Following *in vitro* refolding, a broad peak was observed on HPLC for both proteins (Fig. 2C,D, red trace). The major fraction (indicated by an asterisk) was manually collected, lyophilized, and analyzed by SDS-PAGE (Fig. 2C,D, inner panel). This analysis revealed a prominent monomer band (indicated by an asterisk) along with several faint higher-molecular-weight bands, present with or without dithiothreitol (DTT) treatment, indicating that Dr-CXCL17 and Dr-CXCL17-like are prone to intermolecular cross-linking via isopeptide bonds.

### Activation of the zebrafish GPR25 by the recombinant zebrafish CXCL17s

To determine whether Dr-CXCL17 and Dr-CXCL17-like function as ligands for Dr-GPR25, we used the NanoLuc Binary Technology (NanoBiT)-based β-arrestin recruitment assay, previously validated with Hs-CXCL17 in our recent study [30]. Upon addition of NanoLuc substrate to living HEK293T cells coexpressing the C-terminally large NanoLuc fragment (LgBiT)-fused Dr-GPR25 (Dr-GPR25-LgBiT) and the N-terminally SmBiT tag-fused human β-arrestin 2 (SmBiT-ARRB2), only low baseline bioluminescence was detected (Fig. 3A). Subsequent addition of recombinant 6×His-Dr-CXCL17 led to a rapid, dose-dependent increase in bioluminescence (Fig. 3A), with significant activation observed at concentrations as low as 10 nM. From the dose–response curve (Fig. 3A, inner panel), the EC□□ value for 6×His-Dr-CXCL17 activation of Dr-GPR25 was estimated at approximately 100 nM. In contrast, removal of the three C-terminal residues drastically reduced activity: the truncated 6×His-Dr-□desC3]CXCL17 elicited only minimal bioluminescence increases, even at 1.0 μM (Fig. 3B).

**Fig. 3.**
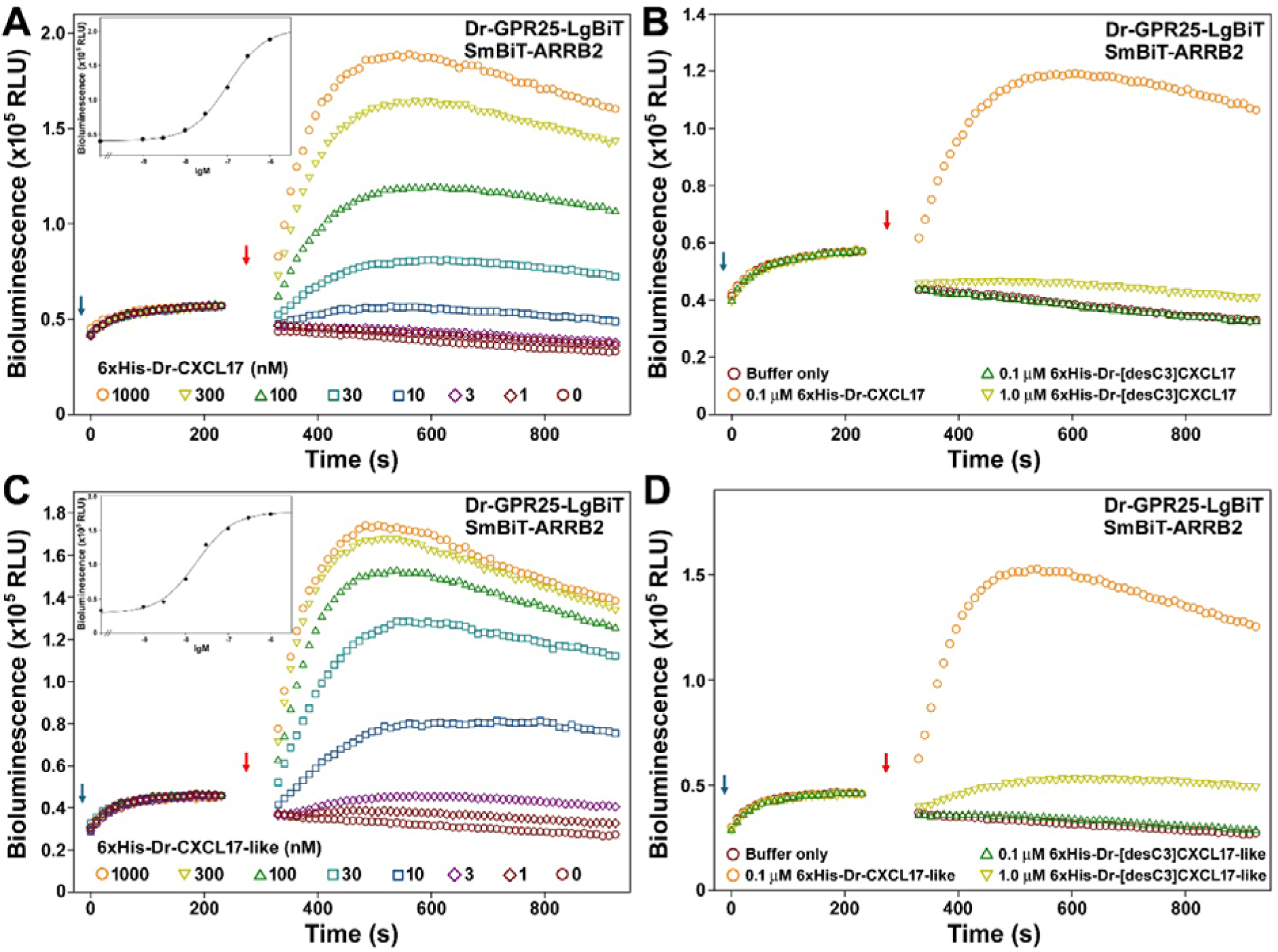
NanoBiT-based β-arrestin recruitment assays of the zebrafish GPR25 induced by recombinant zebrafish CXCL17 or CXCL17-like. (**A,B**) Effect of the WT or truncated zebrafish CXCL17. (**C,D**) Effect of the WT or zebrafish CXCL17-like. In these assays, NanoLuc substrate and different concentrations of WT or truncated zebrafish CXCL17 or CXCL17-like were sequentially added to living HEK293T cells coexpressing Dr-GPR25-LgBiT and SmBiT-ARRB2, and bioluminescence data were measured on a plate reader. Typical bioluminescence change profiles are shown in panels A□D, and the calculated dose response curves of the WT proteins are shown in inner panels of panel A and C. The blue arrows indicate addition of NanoLuc substrate, and red arrows indicate addition of the zebrafish CXCL17 or CXCL17-like.

For recombinant 6×His-Dr-CXCL17-like, addition to the NanoBiT-based β-arrestin recruitment assay also produced a rapid increase in bioluminescence (Fig. 3C), with significant activation observed at concentrations as low as 3.0 nM. From the dose-response curve (Fig. 3C, inner panel), the EC□□ value for 6×His-Dr-CXCL17-like activation of Dr-GPR25 was estimated to be approximately 20 nM, indicating that 6×His-Dr-CXCL17-like is more potent than 6×His-Dr-CXCL17. Removal of the three C-terminal residues resulted in a marked loss of activity: the truncated 6×His-Dr-[desC3]CXCL17-like induced only minimal bioluminescence increases, even at 1.0 μM (Fig. 3D).

In summary, the β-arrestin recruitment assay confirmed that both Dr-CXCL17 and Dr-CXCL17-like are potent agonists of Dr-GPR25, indicating that zebrafish GPR25 has two effective endogenous ligands. The three C-terminal residues are critical for the activation of Dr-GPR25 by both proteins, suggesting that Dr-CXCL17 and Dr-CXCL17-like utilize a similar mechanism to engage and activate the receptor.

### Binding of the recombinant zebrafish CXCL17s with the zebrafish GPR25

To assess the direct binding of Dr-CXCL17 and Dr-CXCL17-like to Dr-GPR25, we employed a NanoBiT-based homogeneous binding assay, which is highly specific and minimally influenced by endogenously expressed CXCL17 receptors. This proximity-based assay has been validated for several other GPCRs in recent studies [31□34]. To generate an appropriate tracer for the assay, a low-affinity SmBiT tag was genetically fused to the N-terminus of Dr-CXCL17-like (Fig. S4), as this protein displayed higher activity than Dr-CXCL17 in the β-arrestin recruitment assay. Following bacterial overexpression, purification, and *in vitro* refolding, the resulting 6×His-SmBiT-Dr-CXCL17-like protein retained high activity in the β-arrestin recruitment assay (Fig. 4A), with an EC□□ of approximately 40 nM, indicating that the SmBiT-tagged tracer is capable of binding Dr-GPR25.

**Fig. 4.**
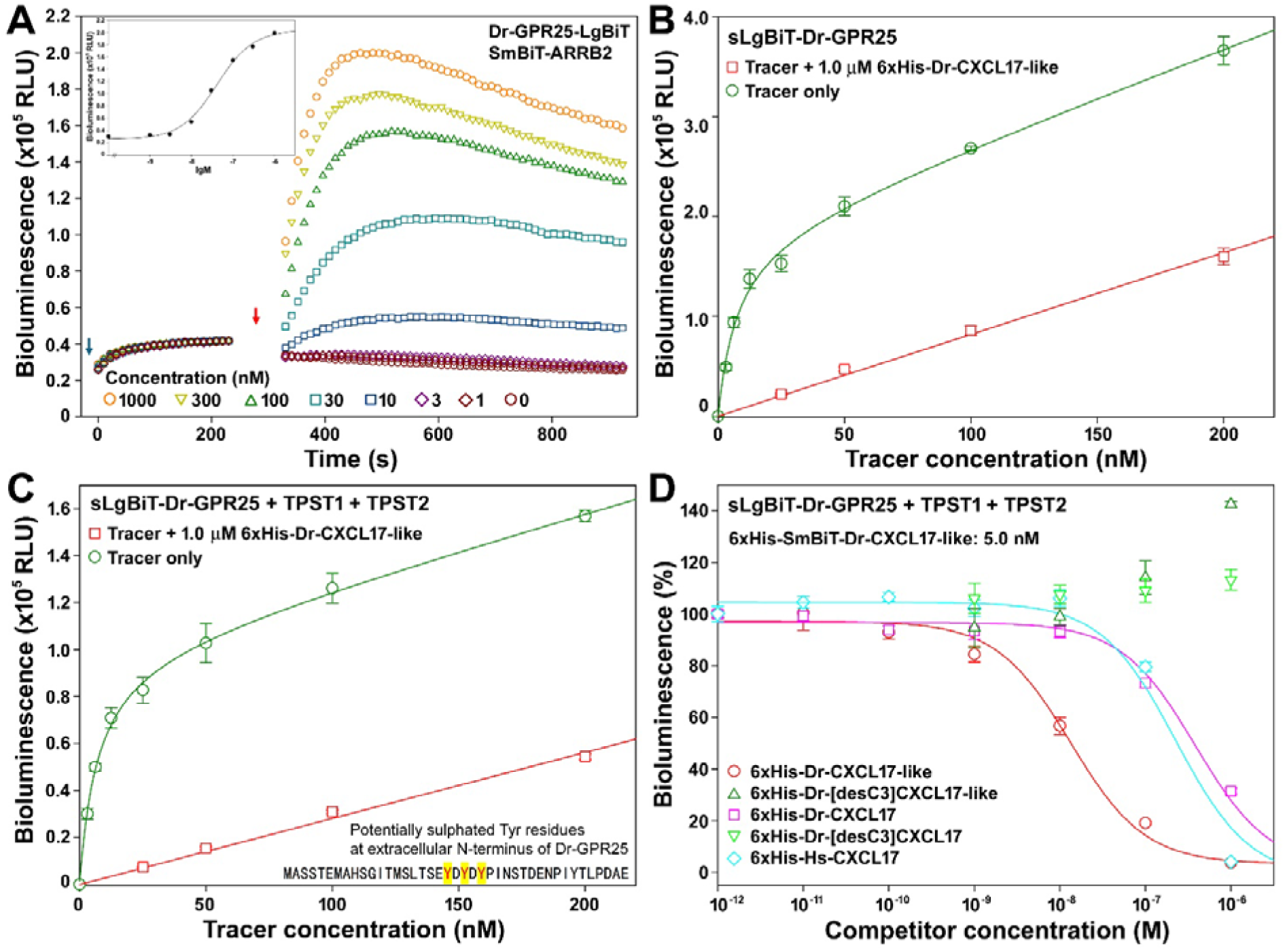
NanoBiT-based binding assays of the zebrafish CXCL17 and CXCL17-like with the zebrafish GPR25. (**A**) Activity of the recombinant 6×His-SmBiT-Dr-CXCL17-like towards Dr-GPR25 measured via the NanoBiT-based β-arrestin recruitment assay. **Inner panel**, dose-response curve of the tracer. The blue arrow indicates the addition of NanoLuc substrate, and the red arrow indicates the addition of peptide. (**B,C**) Saturation binding of 6×His-SmBiT-Dr-CXCL17-like with sLgBiT-Dr-GPR25 without (B) or with (C) coexpression of TPST1 and TPST2. The measured bioluminescence data are expressed as mean ± SD (*n* = 3) and plotted using the SigmaPlot10.0 software. Total binding data (green circles) were fitted with the function of Y = B_max_X/(K_d_+X) + k_non_X, the non-specific binding data (red squares) were fitted with linear curves. (**D**) Competition binding assays of the recombinant zebrafish or human CXCL17s with Dr-GPR25. The measured bioluminescence data are expressed as mean ± SD (*n* = 3) and fitted with sigmoidal curves using the SigmaPlot10.0 software.

We next evaluated the tracer in a saturation binding assay (Fig. 4B). Incubation of recombinant 6×His-SmBiT-Dr-CXCL17-like with living HEK293T cells overexpressing the N-terminally secretory LgBiT (sLgBiT)-fused Dr-GPR25 (sLgBiT-Dr-GPR25) produced a hyperbolic increase in bioluminescence. From the resulting binding curve, the dissociation constant (K_d_) was determined to be 7.5 ± 1.1 nM (*n* = 3). Addition of 1.0 μM unlabeled 6×His-Dr-CXCL17-like markedly reduced the bioluminescence signal (Fig. 4B), confirming that the tracer binds specifically to Dr-GPR25.

The extracellular N-terminus of Dr-GPR25 contains three tyrosine residues that are potentially sulfated, as they are located adjacent to negatively charged residues (Fig. 4C, inner panel). When sLgBiT-Dr-GPR25 was coexpressed with human tyrosylprotein sulfotransferases TPST1 and TPST2, enzymes responsible for tyrosine sulfation [35], a hyperbolic saturation binding curve was obtained (Fig. 4C), yielding a K_d_ value of 7.1 ± 0.9 nM. Coexpression of TPST1 and TPST2 did not significantly alter the K_d_ values, but it slightly reduced the proportion of nonspecific binding (approximately 15% without coexpression vs. around 10% with coexpression at a tracer concentration of 25 nM), suggesting a modest beneficial effect of tyrosylprotein sulfotransferase coexpression on the NanoBiT-based binding assay.

Finally, we performed NanoBiT-based competition binding assays to evaluate the binding potencies of various ligands to Dr-GPR25 (Fig. 4D). Recombinant 6×His-Dr-CXCL17-like efficiently displaced the tracer, resulting in a sigmoidal decrease in bioluminescence, with an IC□□ value of approximately 13 nM. In contrast, the recombinant 6×His-Dr-[desC3]CXCL17-like, lacking the three C-terminal residues, was unable to displace the tracer (Fig. 4D), indicating a complete loss of binding to Dr-GPR25. Interestingly, at high concentrations, this mutant caused a slight increase in bioluminescence (Fig. 5D). This unexpected effect may result from recruitment of additional tracers to the cell surface by the inactive peptide through interactions with both the tracer and cell surface glycosaminoglycans. Consistent with this explanation, human CXCL17, which is positively charged, has been reported to bind strongly to negatively charged cell surface glycosaminoglycans [36].

**Fig. 5.**
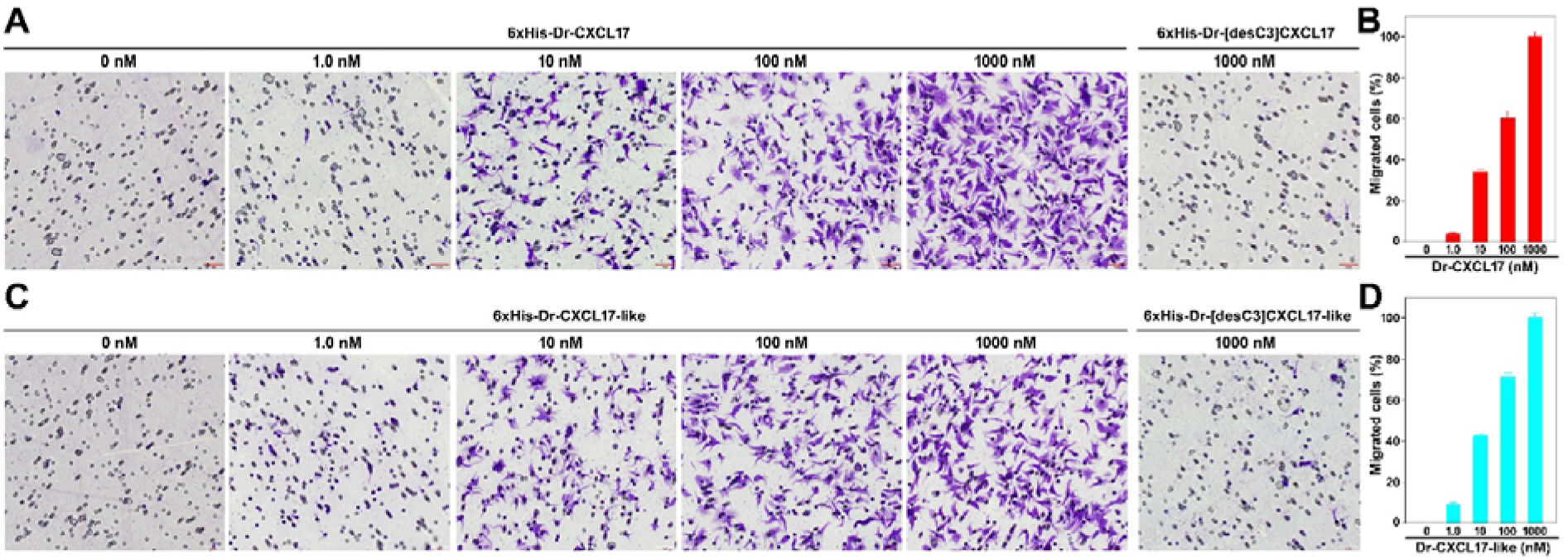
Chemotactic effects of the recombinant zebrafish CXCL17 and CXCL17-like. (**A,B**) Representative images (A) and quantitative analysis (B) of the Dr-GPR25-expressing HEK293T cells after induced by WT or truncated Dr-CXCL17 in the transwell assays. (**C,D**) Representative images (C) and quantitative analysis (D) of the Dr-GPR25-expressing HEK293T cells after induced by WT or truncated Dr-CXCL17-like in the transwell assays. Transfected HEK293T cells were induced to express Dr-GPR25 by Dox, seeded into the permeable membrane-coated inserts, and attracted by chemotactic solution in the lower chamber. After the assay, cells on the upper face of the permeable membrane were wiped off, and cells on the lower face of the permeable membrane were fixed, stained, and observed under a microscope. Representative images of the migrated cells are shown in panel A and C. The scale bar in these images is 50 μm. The migrated cells were quantitatively analyzed using the ImageJ software and the results are expressed as mean ± SD (*n* = 3) and shown in panel B and D.

Recombinant 6×His-Dr-CXCL17 was also able to displace the tracer in the competition binding assay, with an IC□□ of approximately 380 nM (Fig. 4D). This binding potency is about 30-fold lower than that of 6×His-Dr-CXCL17-like, consistent with its reduced activity in the β-arrestin recruitment assay. Removal of the three C-terminal residues abolished binding, as the truncated 6×His-Dr-[desC3]CXCL17 showed no detectable interaction with Dr-GPR25 (Fig. 4D), in agreement with its poor activity in the β-arrestin assay. Recombinant human CXCL17 (6×His-Hs-CXCL17) bound to Dr-GPR25 with an IC□□ of ∼230 nM, comparable to that of 6×His-Dr-CXCL17, consistent with its previously reported high activity toward Dr-GPR25 in the β-arrestin recruitment assay [30].

### Chemotactic activity of the recombinant zebrafish CXCL17s

To evaluate the chemotactic activity of the zebrafish CXCL17s, we performed transwell chemotaxis assays using transiently transfected HEK293T cells expressing Dr-GPR25. As shown in Fig. 5, both recombinant 6×His-Dr-CXCL17 and 6×His-Dr-CXCL17-like induced dose-dependent chemotactic migration of doxycycline (Dox)-induced HEK293T cells, with significant effects observed at concentrations as low as 10 nM. In contrast, the C-terminally truncated mutants of both proteins exhibited almost no chemotactic activity, even at concentrations up to 1.0 μM (Fig. 5), consistent with their reduced activity in receptor activation and binding assays. These findings indicate that both Dr-CXCL17 and Dr-CXCL17-like promote chemotactic migration of transfected HEK293T cells by binding to and activating Dr-GPR25 via their conserved C-terminal fragment.

## Discussion

In this study, we identified two zebrafish CXCL17 paralogs and demonstrated that they are potent agonists of zebrafish GPR25, as confirmed by NanoBiT-based binding and activation assays as well as chemotaxis assays. Owing to their lack of significant amino acid sequence similarity with known mammalian CXCL17s, both paralogs had previously been uncharacterized and their functions unknown. With the rapid advances in DNA sequencing technologies, vast numbers of novel proteins have been discovered across diverse species. The initial step in annotating these proteins is typically to classify them into established protein families based on amino acid sequence similarity with known proteins. However, this approach is ineffective for highly variable proteins, such as the zebrafish CXCL17s reported here, which show no significant overall sequence similarity to functionally characterized counterparts. Consequently, developing strategies to classify and annotate such highly divergent proteins represents an important challenge in the post-genomic era.

Based on three key features derived from mammalian CXCL17s, zebrafish CXCL17s were identified: (1) a C-terminal Xaa-Pro-Yaa motif, where Xaa and Yaa are typically large aliphatic residues; (2) classification as secretory proteins, indicated by an N-terminal signal peptide, with fewer than 200 amino acids; and (3) the presence of six cysteine residues in the mature peptide. These criteria may facilitate the recognition of additional CXCL17 orthologs from other fish species or non-mammalian vertebrates in the future. When combined with amino acid sequence BLAST searches, this approach could enable the identification of more CXCL17 candidates from existing sequence databases. Given that GPR25 orthologs are widely distributed from fishes to mammals, it is reasonable that CXCL17 orthologs are also broadly present among extant vertebrates and remain to be discovered.

CXCL17 has also been reported as an agonist of the MAS-related receptor MRGPRX2 [26]. Our recent findings indicate that human CXCL17 not only activates human MRGPRX2 but also engages the MAS-related receptors MRGPRX1 and MAS1, albeit with slightly lower potency [37]. Notably, activation of these MAS-related receptors by human CXCL17 occurs independently of its conserved C-terminal fragment [37]. In contrast, this conserved C-terminal fragment is essential for CXCL17 orthologs, whether from human or zebrafish, to activate GPR25 orthologs, as demonstrated in the present and previous studies [27,28,30]. These observations suggest that CXCL17 utilizes distinct mechanisms to activate GPR25 and MAS-related receptors. According to NCBI gene database records, GPR25 is broadly distributed from fishes to mammals, whereas MAS-related receptors are restricted to mammals. The identification of CXCL17 orthologs in fishes in this study supports the view that GPR25, rather than MAS-related receptors, represents the evolutionarily conserved receptor for CXCL17. The conserved CXCL17–GPR25 pair likely originated in ancient fishes and subsequently spread across all vertebrate lineages. Our findings provide a foundation for further identification and functional characterization of CXCL17 orthologs in fishes and other non-mammalian vertebrates.

To our knowledge, mammals possess only a single CXCL17 paralog, whereas in this study we identified two CXCL17 paralogs in zebrafish and several other closely related ray-finned fish species. The zebrafish *cxcl17* gene is located near *cnfn*, *pafah1b3*, and *ceacam1* (Fig. S2), a genomic arrangement similar to that of the human *CXCL17* gene and its neighboring genes (Fig. S5). This conservation suggests that CXCL17 orthologs in extant fishes and mammals likely evolved from a common ancestor. In contrast, the zebrafish *cxcl17-like* gene is located on a different chromosome and is flanked by *ponzr3*, *ponzr4*, *ponzr5*, *ponzr6*, *ugt5g1*, and *cldn7a* (Fig. S3), all fish-specific genes, indicating that CXCL17-like may have an independent origin and be restricted to certain fish species. The evolutionary relationship and physiological roles of fish CXCL17 and CXCL17-like warrant further investigation.

## Materials and methods

### Preparation of the recombinant zebrafish CXCL17s

The DNA fragments encoding the mature peptide of Dr-CXCL17 or Dr-CXCL17-like were chemically synthesized using the *E. coli*-biased codons and ligated into a pET vector via Gibson assembly. The resultant expression constructs pET/6×His-Dr-CXCL17 and pET/6×His-Dr-CXCL17-like encode the N-terminally 6×His-tagged zebrafish CXCL17 or CXCL17-like, respectively (Fig. S4). The expression constructs for the C-terminally truncated mutants were generated via the QuikChange approach using pET/6×His-Dr-CXCL17 or pET/6×His-Dr-CXCL17-like as the mutagenesis template. The coding region of the N-terminally 6×His-SmBiT-fused Dr-CXCL17-like was amplified via polymerase chain reaction (PCR) using pET/6×His-Dr-CXCL17-like as the template and then ligated into a pET vector via Gibson assembly, resulting in the expression construct pET/6×His-SmBiT-Dr-CXCL17-like (Fig. S4). The coding regions of Dr-CXCL17 or Dr-CXCL17-like in these expression constructs were confirmed by DNA sequencing.

Overexpression, purification, and refolding of the zebrafish CXCL17s were conducted according to our recent procedure developed for Hs-CXCL17 [28]. Briefly, the transformed bacteria were induced by isopropyl-β-D-thiogalactopyranoside (IPTG) and then collected via centrifugation and lysed by sonication. The overexpressed 6×His-Dr-CXCL17 and 6×His-Dr-CXCL17-like proteins were solubilized from inclusion bodies via an *S*-sulfonation approach, purified by an immobilized metal ion affinity chromatography, and subjected to *in vitro* refolding. The refolded proteins were further purified by HPLC using a C_8_ reverse-phase column (Zorbax 300SB-C8, 9.4 × 250 mm, Agilent Technologies, Santa Clara, CA, USA). The eluted fractions from the reverse-phase column were manually collected, lyophilized, and dissolved in 1.0 mM aqueous hydrochloride (pH 3.0) for quantification and subsequent activity assays. The wild-type (WT) or truncated 6×His-Dr-CXCL17-like and 6×His-SmBiT-Dr-CXCL17-like were quantified by ultra-violet absorbance at 280 nm using the extinction coefficients (ε_280 nm_) of 7450 M^-1^ cm^-1^ and 8940 M^-1^ cm^-1^, respectively. The WT or truncated 6×His-Dr-CXCL17 were quantified by SDS-PAGE using 6×His-Dr-CXCL17-like as standard because they have no ultra-violet absorbance at 280 nm. The samples at different preparation steps were also analyzed by SDS-PAGE.

### Generation of expression constructs for the zebrafish GPR25

The expression constructs for Dr-GPR25 were generated in our recent study [30]. The construct pTRE3G-BI/Dr-GPR25-LgBiT:SmBiT-ARRB2 coexpresses LgBiT-Dr-GPR25 and SmBiT-ARRB2 controlled by a Dox-response bidirectional promoter. The constructs PB-TRE/Dr-GPR25 and PB-TRE/sLgBiT-Dr-GPR25 express an untagged Dr-GPR25 or an N-terminally sLgBiT-fused Dr-GPR25 under control of a Dox-response promoter, respectively.

### The NanoBiT-based **β**-arrestin recruitment assays

The NanoBiT-based β-arrestin recruitment assays were conducted according to our previous procedure [28,30]. Briefly, HEK293T cells were cotransfected with the expression construct pTRE3G-BI/Dr-GPR25-LgBiT:SmBiT-ARRB2 and the expression control vector pCMV-Tet3G (Clontech, Mountain View, CA, USA) using the transfection reagent Lipo8000 (Beyotime Technology, Shanghai, China). Next day, the transfected cells were trypsinized, seeded into white opaque 96-well plates, and cultured in complete Dulbecco’s modification of Eagle’s medium (DMEM) containing 1.0 ng/mL of Dox for ∼24 h to ∼90% confluence. To conduct the β-arrestin recruitment assays, the induction medium was removed, and pre-warmed activation solution (serum-free DMEM plus 1% bovine serum albumin) containing NanoLuc substrate was added (40 μL/well, containing 0.5 μL of NanoLuc substrate stock from Promega, Madison, WI, USA). Thereafter, bioluminescence data were immediately collected for ∼4 min on a SpectraMax iD3 plate reader (Molecular Devices, Sunnyvale, CA, USA). Subsequently, the recombinant WT or truncated zebrafish CXCL17 or CXCL17-like protein (diluted in the activation solution) was added (10 μL/well), and bioluminescence data were continuously collected for ∼10 min. The measured absolute bioluminescence signals were corrected for inter well variability by forcing all curves after addition of NanoLuc substrate to same level and plotted using the SigmaPlot 10.0 software (SYSTAT software, Chicago, IL, USA). To obtain the dose-response curve, the measured bioluminescence data at highest point were plotted with the agonist concentrations using the SigmaPlot 10.0 software (SYSTAT software).

### The NanoBiT-based ligand**□**receptor binding assays

The NanoBiT-based homogenous binding assays were developed using 6×His-SmBiT-Dr-CXCL17-like as a tracer according to our previous procedures for some other GPCRs [31□33]. Briefly, HEK293T cells were transfected with the expression construct PB-TRE/sLgBiT-Dr-GPR25 with or without cotransfection with the tyrosylprotein sulfotransferase expression construct pTRE3G-BI/TPST1:TPST2. Next day, the transfected cells were trypsinized, seeded into white opaque 96-well plates, and cultured in complete DMEM containing 20 ng/mL of Dox for ∼24 h to ∼90% confluence. To conduct the binding assays, the induction medium was removed, and pre-warmed binding solution (serum-free DMEM plus 0.1% bovine serum albumin and 0.01% Tween-20) was added (50 μL/well). For saturation binding assays, the binding solution contains varied concentrations of 6×His-SmBiT-Dr-CXCL17-like. For competition binding assays, the binding solution contains a constant concentration of 6×His-SmBiT-Dr-CXCL17-like and varied concentrations of competitors. To measure cell surface expression level of sLgBiT-Dr-GPR25, binding solution contains 80 nM of synthetic HiBiT peptide. After incubation at room temperature for ∼1 h, diluted NanoLuc substrate (30-fold dilution in the binding solution) was added (10 μL/well), and bioluminescence was immediately measured on a SpectraMax iD3 plate reader (Molecular Devices). The measured bioluminescence data were expressed as mean ± standard deviation (SD, *n* = 3) and plotted using the SigmaPlot 10.0 software (SYSTAT software).

### Chemotaxis assays

The chemotaxis assays were conducted using a transwell apparatus according to our previous procedure [28,30]. Briefly, HEK293T cells were transiently transfected with the expression construct PB-TRE/Dr-GPR25 using the transfection reagent Lipo8000 (Beyotime Technology). Next day, the transfected cells were changed to the complete DMEM containing 1.0 ng/mL of Dox and continuously cultured for ∼24 h. Thereafter, the cells were trypsinized, suspended in serum-free DMEM at the density of ∼5×10^5^ cells/mL, and seeded into polyethylene terephthalate membrane (8 μm pore size)-coated permeable transwell inserts that were pretreated with serum-free DMEM (200 μL/well). The inserts were then put into a 24-well plate containing chemotactic agent (WT or truncated Dr-CXCL17 or Dr-CXCL17-like diluted in serum-free DMEM plus 0.2% bovine serum albumin, 500 μL/well). After cultured at 37°C for ∼5 h, solution in the inserts were removed and cells on the upper face of the permeable membrane were wiped off using cotton swaps, and cells on the lower face of the permeable membrane were fixed with 4% paraformaldehyde solution, stained with crystal violet staining solution (Beyotime Technology), and observed under an Olympus APX100 microscope (Tokyo, Japan). The migrated cells were quantitatively analyzed using the ImageJ software and the results are expressed as mean ± SD (*n* = 3).

## Data availability statement

The data of this study are available in this manuscript, as well as the associated supplementary information.

## Conflicts of Interest

The authors confirm that there is no conflict of interest related to the manuscript.

## Funding

This work was supported by grants from the National Natural Science Foundation of China (31971193, 31470767).

## CRediT Author Contribution

Conceptualization, Z.Y.G.; investigation, J.Y., W.F.H., and J.J.W.; writing – original draft, Z.Y.G.; writing – review & editing, Z.Y.G., and J.Y.; formal analysis, Y.L.L., and Z.G.X.; project administration, Y.L.L., and Z.G.X.; funding acquisition, Z.Y.G.; supervision, Z.Y.G. All authors read and approved the final manuscript.

## Abbreviations

ARRB2: human β-arrestin 2
CXCL17: C-X-C motif chemokine ligand 17
DMEM: Dulbecco’s modification of Eagle’s medium
Dox: doxycycline
DTT: dithiothreitol
GPCR: G protein-coupled receptor
GPR25: G protein-coupled receptor 25
HEK: human embryonic kidney
HiBiT: high-affinity complementation tag for NanoBiT
HPLC: high performance liquid chromatography
IPTG: isopropyl-β-D-thiogalactopyranoside
LgBiT: large NanoLuc fragment for NanoBiT
NanoBiT: NanoLuc Binary Technology
NanoLuc: nanoluciferase
NCBI: the National Center for Biotechnology Information
PCR: polymerase chain reaction
SD: standard deviation
SDS-PAGE: sodium dodecyl sulfate-polyacrylamide gel electrophoresis
sLgBiT: secretory LgBiT
SmBiT: low-affinity complementation tag for NanoBiT
WT: wild-type.

## Notes

### Competing Interest Statement

The authors have declared no competing interest.

