## Supplemental Table S1-S4 and Fig. S1-S5 for "Identification and functional characterization of two CXCL17 paralogs from zebrafish"

**Contents:**

**Table S1.** Information about mammalian CXCL17 orthologs aligned in Fig. S2.

**Table S2.** Information about fish CXCL17 homologs identified via sequence blast with Dr-CXCL17.

**Table S3.** Information about fish CXCL17-like homologs identified via sequence blast with Dr-CXCL17-like.

**Table S4.** Summary of the possible interactions of the zebrafish CXCL17 and CXCL17-like with the zebrafish GPR25 according to the AlphaFold3-predicted structures.

**Fig. S1.** Amino acid sequence alignment of mammalian CXCL17 orthologs.

**Fig. S2.** Position of the zebrafish *cxcl17* gene (*zgc:158701*) in the genome of *Danio rerio*.

**Fig. S3.** Position of the zebrafish *cxcl17-like* gene (*si:dkey-112a7.5*) in the genome of *Danio rerio*.

**Fig. S4.** The nucleotide and amino acid sequence of the zebrafish CXCL17 and CXCL17-like overexpressed in *E. coli*.

**Fig. S5.** Position of the human *CXCL17* gene in human genome.

**Table S1.** Information about mammalian CXCL17 orthologs aligned in Fig. S1. The information was downloaded from the NCBI gene database (https://ncbi.nlm.nih.gov/gene).

| Mammalian species | Gene ID | mRNA ID | Protein ID |
|---|---|---|---|
| *Homo sapiens* | 284340 | NM_198477 | NP_940879 |
| *Pan troglodytes* | 741429 | XM_001154726 | XP_001154726 |
| *Pan paniscus* | 100975966 | XM_003811758 | XP_003811806 |
| *Macaca mulatta* | 708108 | XM_001105835 | XP_001105835 |
| *Sapajus apella* | 116530866 | XM_032249326 | XP_032105217 |
| *Suricata suricatta* | 115281288 | XM_029926676 | XP_029782536 |
| *Galeopterus variegatus* | 103586108 | XM_008567252 | XP_008565474 |
| *Mus musculus* | 232983 | NM_153576 | NP_705804 |
| *Rattus norvegicus* | 308436 | NM_001107491 | NP_001100961 |
| *Mesocricetus auratus* | 101826068 | XM_021223580 | XP_021079239 |
| *Octodon degus* | 101573419 | XM_023704517 | XP_023560285 |
| *Arvicola amphibius* | 119821858 | XM_038340966 | XP_038196894 |
| *Arvicanthis niloticus* | 117720566 | XM_034519100 | XP_034374991 |
| *Mastomys coucha* | 116100415 | XM_031384229 | XP_031240089 |
| *Peromyscus leucopus* | 114685351 | XM_028859975 | XP_028715808 |
| *Grammomys surdaster* | 114632694 | XM_028781282 | XP_028637115 |
| *Marmota marmota* | 107151419 | XM_015496800 | XP_015352286 |
| *Fukomys damarensis* | 104861740 | XM_010623332 | XP_010621634 |
| *Chinchilla lanigera* | 102020166 | XM_005412354 | XP_005412411 |
| *Microtus ochrogaster* | 101982135 | XM_005361160 | XP_005361217 |
| *Heterocephalus glaber* | 101708913 | XM_004873008 | XP_004873065 |
| *Cavia porcellus* | 100734695 | XM_013146847 | XP_013002301 |
| *Felis catus* | 101086020 | XM_003997765 | XP_003997814 |
| *Canis lupus familiaris* | 111090090 | XM_038656810 | XP_038512738 |
| *Mustela putorius furo* | 101683082 | XM_004780427 | XP_004780484 |
| *Leopardus geoffroyi* | 123578900 | XM_045442260 | XP_045298216 |
| *Prionailurus bengalensis* | 122493991 | XM_043598820 | XP_043454755 |
| *Panthera leo* | 122207597 | XM_042917645 | XP_042773579 |
| *Puma yagouaroundi* | 121018500 | XM_040457200 | XP_040313134 |
| *Vulpes lagopus* | 121484348 | XM_041743498 | XP_041599432 |
| *Hyaena hyaena* | 120240684 | XM_039245502 | XP_039101433 |
| *Lontra canadensis* | 116857606 | XM_032841857 | XP_032697748 |
| *Mustela erminea* | 116580309 | XM_032326546 | XP_032182437 |
| *Ursus arctos* | 113243523 | XM_026482222 | XP_026338007 |
| *Vulpes vulpes* | 112931475 | XM_026014200 | XP_025869985 |
| *Puma concolor* | 112851386 | XM_025914921 | XP_025770706 |
| *Pteropus vampyrus* | 105307357 | XM_011382839 | XP_011381141 |
| *Panthera tigris* | 102961962 | XM_007086589 | XP_007086651 |
| *Pteropus alecto* | 102889606 | XM_006903850 | XP_006903912 |
| *Lutra lutra* | 125088637 | XM_047709872 | XP_047565828 |
| *Meles meles* | 123931218 | XM_045988111 | XP_045844067 |
| *Neogale vison* | 122912075 | XM_044257394 | XP_044113329 |
| *Talpa occidentalis* | 119249391 | XM_037516694 | XP_037372591 |
| *Sturnira hondurensis* | 119000497 | XM_037065531 | XP_036921426 |
| *Loxodonta africana* | 100668516 | XM_003406663 | XP_003406711 |
| *Bos taurus* | 788717 | NM_001083799 | NP_001077268 |
| *Camelus ferus* | 102511827 | XM_032485980 | XP_032341871 |
| *Odocoileus virginianus* | 110123322 | XM_020871140 | XP_020726799 |
| *Bison bison bison* | 104997641 | XM_010852534 | XP_010850836 |
| *Bubalus bubalis* | 102401266 | XM_006052397 | XP_006052459 |
| *Capra hircus* | 102181231 | XM_018062511 | XP_017918000 |
| *Ochotona princeps* | 101528914 | XM_012930594 | XP_012786048 |
| *Balaenoptera musculus* | 118885645 | XM_036834491 | XP_036690386 |
| *Halichoerus grypus* | 118542831 | XM_036103355 | XP_035959248 |
| *Mirounga leonina* | 117998055 | XM_034986582 | XP_034842473 |
| *Tursiops truncatus* | 117309080 | XM_033845069 | XP_033700960 |
| *Phocoena sinus* | 116744452 | XM_032614237 | XP_032470128 |
| *Phoca vitulina* | 116622190 | XM_032388455 | XP_032244346 |
| *Monodon monoceros* | 114903747 | XM_029236980 | XP_029092813 |
| *Eumetopias jubatus* | 114197075 | XM_028087908 | XP_027943709 |
| *Zalophus californianus* | 113935480 | XM_027617446 | XP_027473247 |
| *Lagenorhynchus obliquidens* | 113605953 | XM_027079714 | XP_026935515 |
| *Callorhinus ursinus* | 112807009 | XM_025849582 | XP_025705367 |

| | | | |
|---|---|---|---|
| *Physeter catodon* | 102983204 | XM_024132473 | XP_023988241 |
| *Trichechus manatus latirostris* | 101348725 | XM_004388765 | XP_004388822 |
| *Orcinus orca* | 101278327 | XM_004271217 | XP_004271265 |
| *Equus caballus* | 100629292 | XM_003362327 | XP_003362375 |
| *Choloepus didactylus* | 119521473 | XM_037819798 | XP_037675726 |
| *Tupaia chinensis* | 102492896 | XM_014582832 | XP_014438318 |
| *Erinaceus europaeus* | 103122352 | XM_007532948 | XP_007533010 |
| *Manis pentadactyla* | 118918052 | XM_036896470 | XP_036752365 |
| *Myotis myotis* | 118674385 | XM_036346119 | XP_036202012 |
| *Desmodus rotundus* | 112320178 | XM_024577629 | XP_024433397 |
| *Pipistrellus kuhlii* | 118702335 | XM_036408200 | XP_036264093 |
| *Molossus molossus* | 118639378 | XM_036275769 | XP_036131662 |
| *Rhinolophus ferrumequinum* | 117035523 | XM_033129434 | XP_032985325 |
| *Phyllostomus discolor* | 114511635 | XM_028530396 | XP_028386197 |
| *Miniopterus natalensis* | 107534226 | XM_016209250 | XP_016064736 |
| *Rousettus aegyptiacus* | 107502574 | XM_016129904 | XP_015985390 |
| *Phascolarctos cinereus* | 110219727 | XM_021003326 | XP_020858985 |
| *Trichosurus vulpecula* | 118835868 | XM_036743155 | XP_036599050 |
| *Sarcophilus harrisii* | 116422895 | XM_031963247 | XP_031819107 |
| *Dromiciops gliroides* | 122745891 | XM_043991340 | XP_043847275 |
| *Vombatus ursinus* | 114040160 | XM_027858170 | XP_027713971 |
| *Ornithorhynchus anatinus* | 103166888 | XM_029066640 | XP_028922473 |
| *Tachyglossus aculeatus* | 119946135 | XM_038767563 | XP_038623491 |

**Table S2.** Information about fish CXCL17 homologs identified via sequence blast with Dr-CXCL17. The protein sequence blast was conducted via the NCBI online server (https://blast.ncbi.nlm.nih.gov/Blast.cgi), and information about the retrieved homologs was downloaded from the NCBI gene database or nucleotide database.

| Fish species | Gene ID | mRNA ID | Protein ID | Amino acid sequence |
|---|---|---|---|---|
| *Danio rerio* | 100151367 | NM_001144821 | NP_001138293 | MKTMNFQILVLAFAVMIVTNIQCEARPQEGKSDKSAEVKGHAMPRKCNCQVRGTALDRNCVCEMPHKSRPTLNPEQKNMCLKKKIKTFRKCLQFMGANKKIAKGASLPI |
| *Danio aesculapii* | 130242767 | XM_056474668 | XP_056330643 | MNFQILVLAFAVVTVTNLQCQARPQVENSDKSAEVKGHATPRKCNCRGRGTALDQNCVCEMPPKSRQTLSPEQKNMCLKKRTRTFRKCQQLIGANKKSAKAISMPI |
| *Ictalurus furcatus* | 128607944 | XM_053625239 | XP_053481214 | MKFVMLLLVFTAMVGFSALCQAVIDSEVSDKNVTASSGDAVGPADVTSHTRSTKQRGANCACAGKSKAQRGCQCKMNNFKHGLSPEERVLCLKKGIRNYKKCKSVILKTVKTKEDPKQISIPV |
| *Ictalurus furcatus* | 128607944 | XM_053625248 | XP_053481223 | MKFVMLLLVFTAMVGFSALCQVIDSEVSDKNVTASSGDAVGPADVTSHTRSTKQRGANCACAGKSKAQRGCQCKMNNFKHGLSPEERVLCLKKGIRNYKKCKSVILKTVKTKEDPKQISIPV |
| *Ictalurus punctatus* | 108266547 | XM_017470017 | XP_017325506 | MKFVMLLLVFTAMVGFSALCQAVIDSQVSDKNVTASSGDAVGSADVTSHTRSTKQRSANCVCAGRSKARRGCQCKMNNFKYGLSPEERVLCLNKGIRNYKKCKTVILKTVRTKEDPKQISIPM |
| *Ictalurus punctatus* | 108266547 | XM_017470026 | XP_017325515 | MKFVMLLLVFTAMVGFSALCQVIDSQVSDKNVTASSGDAVGSADVTSHTRSTKQRSANCVCAGRSKARRGCQCKMNNFKYGLSPEERVLCLNKGIRNYKKCKTVILKTVRTKEDPKQISIPM |
| *Labeo rohita* | 127178215 | XM_051130938 | XP_050986895 | MNFQILLLAFAVVIATNIHCQAAQPQLRDSDKSPFPKGLAIFKQQGKKCNCLGIRDPLNQNCHCEMQRQLRMKEQKTLCLKNGILSKKCLELAGGNRKERKGFSMPI |
| *Labeo rohita* | 127178215 | XM_051130939 | XP_050986896 | MNFQILLLAFAVVIATNIHCQAQPQLRDSDKSPFPKGLAIFKQQGKKCNCLGIRDPLNQNCHCEMQRQLRMKEQKTLCLKNGILSKKCLELAGGNRKERKGFSMPI |
| *Pseudorasbora parva* | 137083191 | XM_067448952 | XP_067305053 | MKVQVILLAFAVVVATNIHCQVQSQLTDADTSPVFKGHVMSRQVWKCNCNGRRISLEQNCPCELQRQNRMLSKEQRAVCQKKGIVTYKKCQQLTGGNRKEKNVRKVYSMPI |
| *Carassius carassius* | 132158314 | XM_059567675 | XP_059423658 | MNFQVPLLVFAVVITTSINCQEQPQVGESNKSPVVKGQVISRHQGKTCNCIGKRNALEQNCPCGLQRQNRILSNDKKAWCQKKGISTFKKCQQLIGENRKEKKGISMPI |
| *Onychostoma macrolepis* | 131522610 | XM_058748243 | XP_058604226 | MNFQLLLLVFAVVIATNIHCQAQPPLGNSNKSPAVKGQVISRQQGKTCNCTGRRNALEQNNCPCELQRQYRILSKEQKALCLKKGISTFKKCQQLIGGNRKEKKGFSMPI |
| *Carassius gibelio* | 128030755 | XM_052618693 | XP_052474653 | MRNFQVLLLVFAVVIATNIHIQAKPQLGDSDKSPVVKSKQQSKPCSCVGRRNALEQDNCTCERQRQHGTLSEEQRTCQKKKYRKCPRLNRGNKKEEKGISMPI |
| *Myxocyprinus asiaticus* | 127452759 | XM_051718450 | XP_051574410 | MKLQVLLLAFAVFIASVHCQAQAQLGQSHKSPVIKGQVLLSRQQGRSCNCSGRRNGLELNCPCELHRQHRVLSQKQWALCQKKLIQTYKKCQQMIFGEKRKEKKGNKGISNPI |
| *Megalobrama amblycephala* | 125243866 | XM_048153711 | XP_048009668 | MKFHMLLLAFVVVIVTNIHCQVQPQLEDSDKSPVDKGHVMSKRQVRTCNCGGRRNALEQNCPCERQRQYEILSKEQRAFCQKKGIWTYKKCQQMIGGNRKEKKGNKGFSMPI |
| *Puntigrus tetrazona* | 122360472 | XM_043261100 | XP_043117035 | MNFQVLLLVCAVVIATNIHCQAQPQLGDSNKSPVVKGHIISVQQGKKCNCIGRRNALEQNYCPCEQQRQYKLLSKEQTFACLKKGISTFKKCQQFTGGNKKKKVISTPF |
| *Pimephales promelas* | 120484472 | XM_039679646 | XP_039535580 | MKFQVLLLAFALVIATNIYCQAQPQRRDSDKSPEVSRRQVRTCNCSGRRFENCPCELQRQYRVLKEQKAFCQKKGSATSKICQRLTGGIRKQKKGNIGSMPI |
| *Paramisgurnus dabryanus* | 141280008 | XM_073811699 | XP_073667800 | MKFQILLLTCAVLIATDVYGEAQPQQGDSDKSMSLQQGRPCNCIERRNGQKQNCPCSLPSQRTILSEKQRDLCKKKGIKTFKKCKQLIPQNIKNGKKSNKAMGTPF |
| *Cirrhinus molitorella* | | KAL1259315 | QQF64_009892 | MMNFQVLLLAFAVVIATNIHCQALPQLGDSNKSPVVEGLTISRQPSKLCNCIGRRNALDQNYCTCEMQRQHRMLNKEQIALCLKKGISTYKKCLQWTGGNRKDRKAISMPI |
| *Cirrhinus molitorella* | | KAK2903191 | Q8A67_007904 | MNFQVLLLAFAVVIATNIHCQALPQLGDSNKSPVVEGLTISRQPSKLCNCIGRKNALDQNYCTCEMQRQHRMLNKEQIALCLKKGISTYKKCLQLTGGNRKDRKAISMPI |
| *Culter alburnus* | | KAK9960771 | ABG768_008606 | MKFQVLLLAFVVVIVTNIHCQVQPQLEDSDKSPVDKGHVMSKRQVRTCNCGGRRNALEQNCPCELQRQYKILSKEQRAFCQKKGIGTYKKCQQMIGGNRKEKKGNKGFSMPI |
| *Anabarilius grahami* | | ROI36443 | DPX16_11384 | MKFQVILLAFAVVIVTNIHCQAQHQLEDSDKSPVDKGHIMSKRQVRTCNCRGKRNALEQNCPCELQRQYGILSKEQKAFCQKKGIGTYKKCQQRTGGNRKEKKGNKGFSMPI |
| *Leuciscus waleckii* | | XDV40076 | PO909_009229 | MKFQMLLLAFALVIATNIHCQAQPQLRDSDKSPEVKGHVMSRRQVRTCNCSGRRISVEQNCPCAQRLLSKEQRALCQKKGIASFKKCQQLNGGIRKQKKGNISMPI |
| *Cyprinus carpio* | | KTF95749 | cypCar_00039330 | MNFQVLLLVFAVVIATNIHCQAQPRLGDSDKSPVDKSKQQGKTCNCTGRQNALEQNNCPCESQYRILSKERAVNERALCQKKGILTKKCQHMNRSNRKEKKGISMPI |
| *Phoxinus phoxinus* | | KAK7147560 | R3I94_010169 | MKFQVLLLAFALVIATNIHCQAQPQLRDSDKSPEGNGYIMSRRQVRTCNCNGFLSKEKKAFCQKKGIATSKKCKQLTGGNRKQMKANNGPSMP |

**Table S3.** Information about fish CXCL17-like homologs identified via sequence blast with Dr-CXCL17-like. The protein sequence blast was conducted via the NCBI online server (https://blast.ncbi.nlm.nih.gov/Blast.cgi), and information about the retrieved homologs was downloaded from the NCBI gene database or nucleotide database.

| Fish species | Gene ID | mRNA ID | Protein ID | Amino acid sequence |
|---|---|---|---|---|
| *Danio rerio* | 100536854 | NM_001386806<br>XM_073906074 | NP_001373735<br>XP_073762175 | MTKPICLVFALLILTIILCNNSVCSQRRSMKQSAVCGCKLYPDKGLKCTKRPNPKSRDE<br>YYEILKCICRDTQIFSKSSRKEYLKRCNKFYPSLPL |
| *Misgurnus anguillicaudatus* | 129439872 | XM_073859897 | XP_073715998 | MSKTIYLVFVLVILTTLLCNSPVCSQRGSGNGQATCGCKIHPRKGLQCVRRPHTTWAEI<br>VQCICNNRKTYLNGDSKRLYQKYCNRTISTPL |
| *Chaetodon trifascialis* | 139351666 | XM_070993653 | XP_070849754 | MSRIIVVSLLLIILVDNFYHSTASHAEFRVSKVHIRKCRCRVFPDGRIKCRSPLLPSNY<br>VKKQDLIKCFCNKSNQHKFPEFQAACGSWRPNIPITLL |
| *Garra rufa* | 141331768 | XM_073836795 | XP_073692896 | MSKPICLLFVLVILTTILCYNPVCSQRGSLKQKTACGCKIHPNGSLWCAKRHNPKNSYE<br>YDEVVRCICRNPTKYLNENSKKQFVRMCHSKYPSLPL |
| *Oncorhynchus clarkii lewisi* | 139417144 | XM_071166389 | XP_071022490 | MSKLICVALLLVFLVSIWCHNTVSSTKWSVRDVLRKCRCKVLPNGREICRKPLFPKTPE<br>ETKQLIKCFCRKYKLYKHSAKLNLQTKRDLEKCSFIWSNPF |
| *Oncorhynchus mykiss* | 110531667 | XM_021614986 | XP_021470661 | MSKLICVALLLVFLVSIWCHNTVSSTKWSVRDVLRKCRCKVLPNGREICRKPLFPKTPE<br>ETKQLIKCFCRKYKLYKHSAKLNLETKRDLEKCSFIWSNPF |
| *Oncorhynchus keta* | 118368844 | XM_035753275 | XP_035609168 | MSKLICVALLLVFLVSIWCHNTVSSTKWSVRDVLRKCRCKVLPNGREICRKPLFPKTPE<br>ETKQLIKCFCRKYNLYKHSAKLNLETKRDLEKCSFIWSNPF |
| *Oncorhynchus kisutch* | 109906877 | XM_031791963 | XP_031647823 | MSKLICVALLLVFLVSIWCHNTVSSTKWSVRDVLRKCRCKVLPNGREICRKPLFPKTPE<br>ETKQLIKCFCRKYNLYKHGAKLNLETKRDLEKCSFIWSNPF |
| *Leuciscus waleckii* | | XDV19005 | PO909_024587 | MSKPICLVFALLILTTILCNNSVCSQRGSWKQSAVCGCKIHHDNSLRCTKKHSPKTLEE<br>YHKMVGCICSDRQKYFNSKKQFLNMCKSYSQTPL |
| *Cirrhinus molitorella* | | KAK2913578 | Q8A67_001977 | MSKPICLLFVLVILTTILCYNPVCSQRGSLKQIAACGCNFHPNGSLRCTKRHNLKTGYE<br>YNEVVKCICRNPTKYLNEDLKKTFVRMCNKLSMPL |
| *Phoxinus phoxinus* | | KAK7171077 | R3I94_001092 | MSKPICLVFALLILTTILCNNSVCSQRGSSKQSAVCGCTIHHNSLRCTKRHRPKTIEEN<br>HKMISCICNNPQKYLNEASKKQFRNMCISNSQTPF |
| *Phoxinus phoxinus* | | KAK7177025 | R3I93_001084 | MSKPICLVFAVLILTTILCNNSVCSQRGSSKQSAVCGCTIHHNSLRCTKRHRPKTIEEN<br>HKMISCICNNPQKYLNEASKKQFRNMCISNSQTPF |
| *Triplophysa rosa* | | KAI7806138 | IRJ41_001095 | MSKTVSLVFVLVILTTILCNNSVCCRGFWKGRISCGCKIHPEKGLQCSKRHNFKTMDEI<br>MKCICRNPRTYLTDSSKRLFQKMCKPNISTPL |
| *Megalops atlanticus* | | KAG7465314 | MATL_G00175080 | MSKLCCGSLLLILLAVIWCDYSVESRRWISPNRLLKDECKCRVLANKKVSCRKDFSPKT<br>MQERFDMLKCLCKKHYNELEMKMDADFQKACKKIFRDVPIPQPLS |

**Table S4.** Summary of the possible interactions of the zebrafish CXCL17 and CXCL17-like with the zebrafish GPR25 according to the AlphaFold3-predicted structures. The AlphaFold3 prediction was conducted via the online server (https://alphafoldserver.com). The receptor residues involving ligand-binding are indicated by asterisks in Fig. 1B.

| Ligand | Ligand residue | Interacting residues in Dr-GPR25 |
|---|---|---|
| Dr-CXCL17 | C-terminal carboxyl moiety | ε-amine moiety of K274 (TMD6) |
| | I109 | I127 (TMD3); F270 (TMD6); F307 (TMD7) |
| | P108 | W104 (TMD2) |
| | L107 | L299 (TMD7); M300 (TMD7) |
| Dr-CXCL17-like | C-terminal carboxyl moiety | ε-amine moiety of K274 (TMD6) |
| | L95 | I127 (TMD3); F270 (TMD6); F307 (TMD7) |
| | P94 | W104 (TMD2) |
| | L93 | L299 (TMD7); M300 (TMD7) |

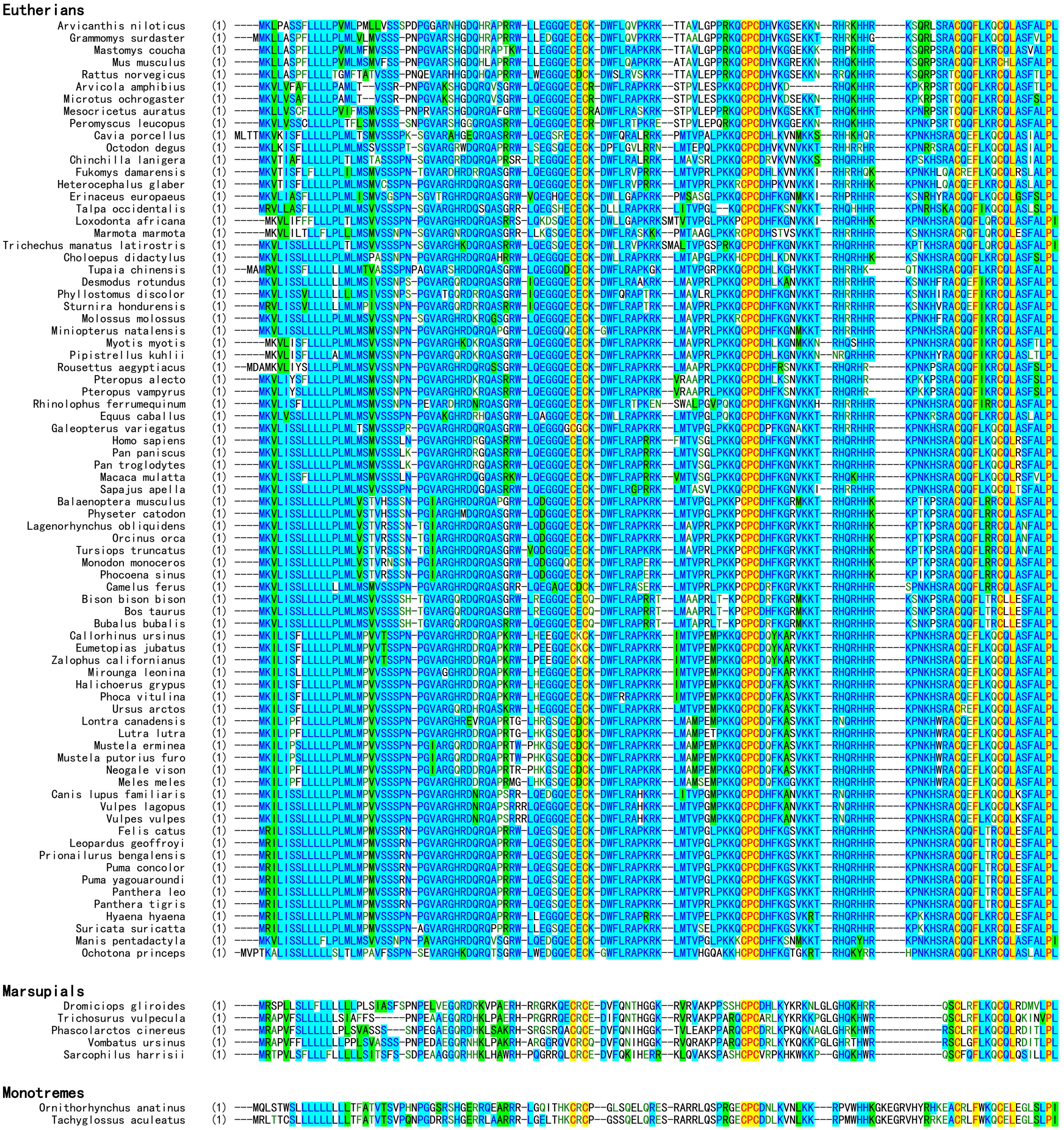


**Fig. S1.** Amino acid sequence alignment of mammalian CXCL17 orthologs. Accession numbers of these orthologs are listed in Table S1. These sequences were aligned via AlignX algorithm using the Vector NTI 11.5.1 software.

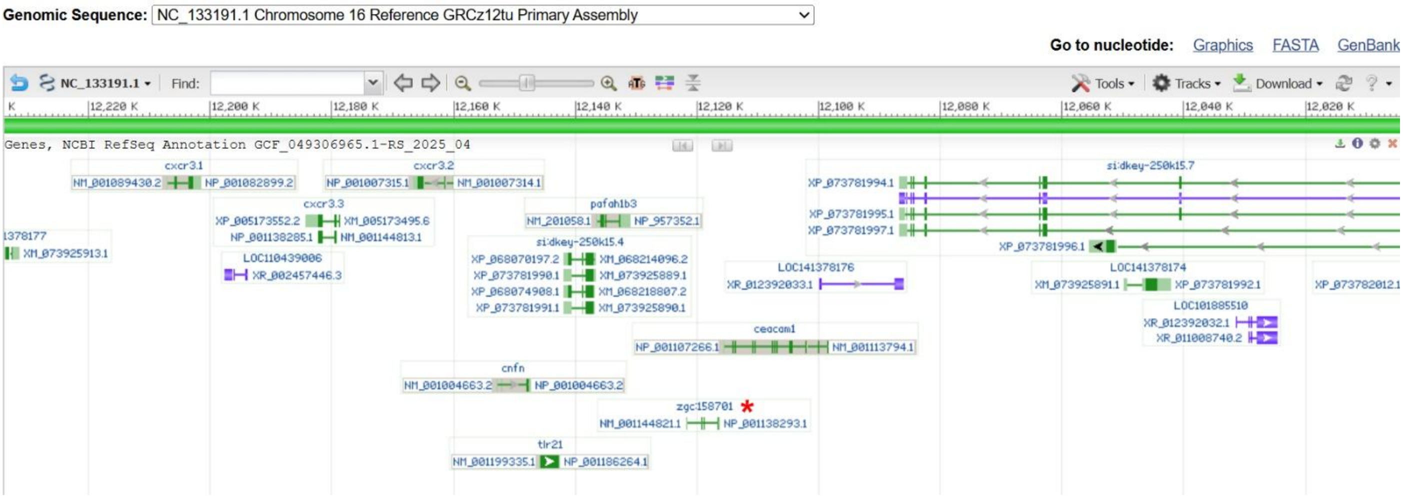


**Fig. S2.** Position of the zebrafish *cxcl17* gene (*zgc:158701*) in the genome of *Danio rerio*. The zebrafish *cxcl17* gene (*zgc:158701*) is indicated by a red asterisk. The information was downloaded from the NCBI gene database (https://www.ncbi.nlm.nih.gov/gene/?term=100151367).

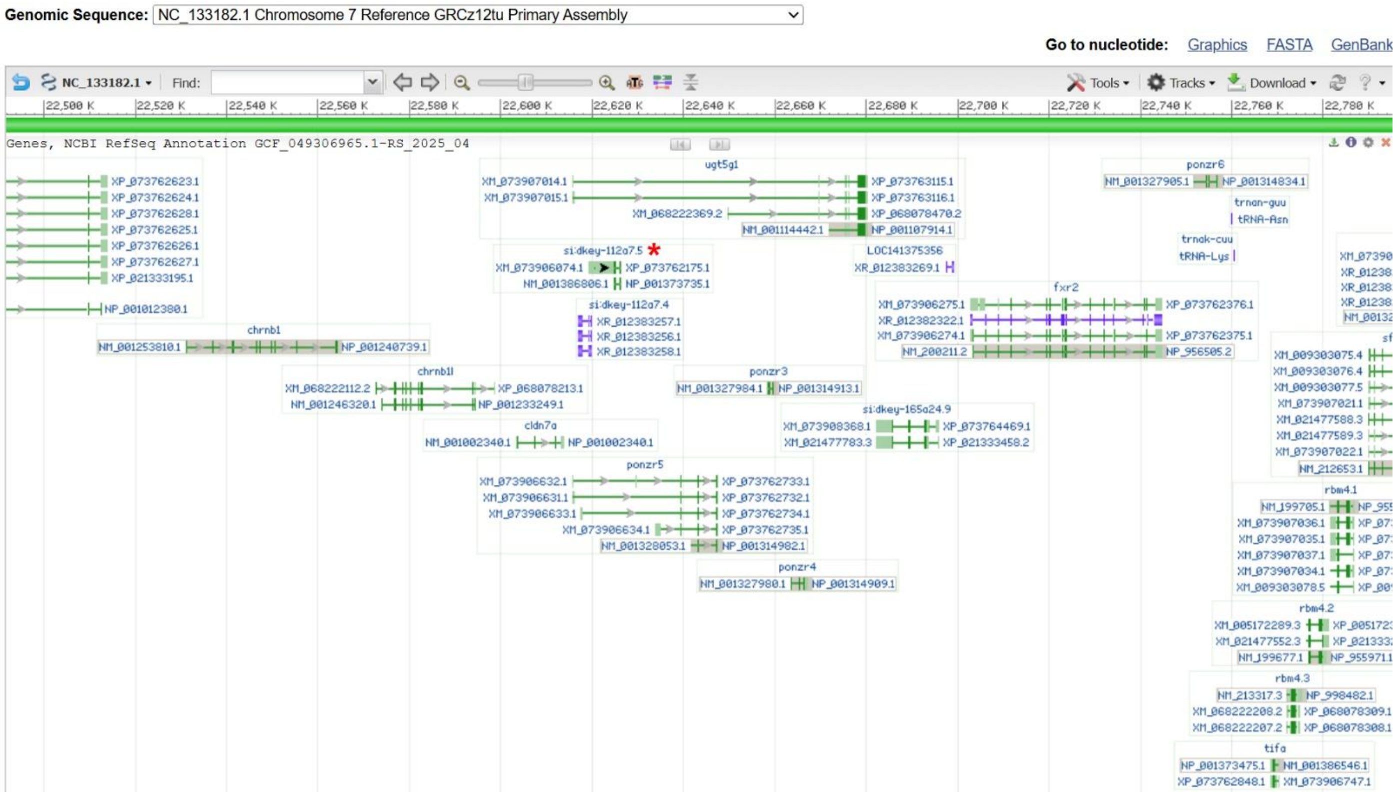


**Fig. S3.** Position of the zebrafish *cxcl17-like* gene (*si:dkey-112a7.5*) in the genome of *Danio rerio*. The zebrafish *cxcl17-like* gene (*si:dkey-112a7.5*) is indicated by a red asterisk. The information was downloaded from the NCBI gene database (https://www.ncbi.nlm.nih.gov/gene/?term=100536854).

### 6×His-Dr-CXCL17

```
1   CTT TAA GAA GGA GAT ATA ATG CAT CAT CAC CAC CAT CAC CGT CCG CAG GAA GGT AAG TCC GAC AAA TCT GCA GAG
    GAA ATT CTT CCT CTA TAT TAC GTA GTA GTG GTG GTA GTG GCA GGC GTC CTT CCA TTC AGG CTG TTT AGA CGT CTC
                             M   H   H   H   H   H   H   R   P   Q   E   G   K   S   D   K   S   A   E

76  GTT AAA GGC CAT GCT ATG CCT CGC AAA TGC AAC TGT CAA GTG CGT GGT ACT GCG CTG GAT CGC AAC TGT GTG TGT
    CAA TTT CCG GTA CGA TAC GGA GCG TTT ACG TTG ACA GTT CAC GCA CCA TGA CGC GAC CTA GCG TTG ACA CAC ACA
     V   K   G   H   A   M   P   R   K   C   N   C   Q   V   R   G   T   A   L   D   R   N   C   V   C

151 GAA ATG CCA CAC AAA AGC CGT CCG ACC CTC AAT CCA GAA CAG AAA AAC ATG TGC TTA AAG AAG AAA ATT AAA ACC
    CTT TAC GGT GTG TTT TCG GCA GGC TGG GAG TTA GGT CTT GTC TTT TTG TAC ACG AAT TTC TTC TTT TAA TTT TGG
     E   M   P   H   K   S   R   P   T   L   N   P   E   Q   K   N   M   C   L   K   K   K   I   K   T

226 TTT CGC AAA TGC CTG CAG TTT ATG GGT GCA AAC AAG AAA ATC GCG AAA GGC GCC AGT TTG CCG ATT TAA GCG GCC
    AAA GCG TTT ACG GAC GTC AAA TAC CCA CGT TTG TTC TTT TAG CGC TTT CCG CGG TCA AAC GGC TAA ATT CGC CGG
     F   R   K   C   L   Q   F   M   G   A   N   K   K   I   A   K   G   A   S   L   P   I   *

301 GCA CTC GAG CAC CAC
    CGT GAG CTC GTG GTG
```

### 6×His-Dr-CXCL17-like

```
1   CTT TAA GAA GGA GAT ATA ATG CAT CAT CAC CAC CAT CAC CAG CGT CGC TCC ATG AAA CAA TCT GCG GTG TGT GGC
    GAA ATT CTT CCT CTA TAT TAC GTA GTA GTG GTG GTA GTG GTC GCA GCG AGG TAC TTT GTT AGA CGC CAC ACA CCG
                             M   H   H   H   H   H   H   Q   R   R   S   M   K   Q   S   A   V   C   G

76  TGC AAA CTT TAC CCT GAT AAG GGT CTG AAA TGT ACC AAA CGT CCG AAT CCA AAA TCG CGC GAT GAA TAC TAT GAG
    ACG TTT GAA ATG GGA CTA TTC CCA GAC TTT ACA TGG TTT GCA GGC TTA GGT TTT AGC GCG CTA CTT ATG ATA CTC
     C   K   L   Y   P   D   K   G   L   K   C   T   K   R   P   N   P   K   S   R   D   E   Y   Y   E

151 ATT CTG AAG TGT ATT TGC CGT GAC ACC CAG ATC TTC TCA AAA AGC AGT CGC AAA GAA TAT CTG AAG CGT TGC AAC
    TAA GAC TTC ACA TAA ACG GCA CTG TGG GTC TAG AAG AGT TTT TCG TCA GCG TTT CTT ATA GAC TTC GCA ACG TTG
     I   L   K   C   I   C   R   D   T   Q   I   F   S   K   S   S   R   K   E   Y   L   K   R   C   N

226 AAA TTT TAC CCG TCT CTC CCG TTG TAA GCG GCC GCA CTC GAG CAC CAC
    TTT AAA ATG GGC AGA GAG GGC AAC ATT CGC CGG CGT GAG CTC GTG GTG
     K   F   Y   P   S   L   P   L   *
```

### 6×His-SmBiT-Dr-CXCL17-like

```
1   CTT TAA GAA GGA GAT ATA ATG CAT CAT CAC CAT CAC CAT GGT GTG ACC GGC TAC CGT CTG TTT GAA GAA ATT CTG
    GAA ATT CTT CCT CTA TAT TAC GTA GTA GTG GTA GTG GTA CCA CAC TGG CCG ATG GCA GAC AAA CTT CTT TAA GAC
                             M   H   H   H   H   H   H   G   V   T   G   Y   R   L   F   E   E   I   L

     G   G   Q   R   R   S   M   K   Q   S   A   V   C   G   C   K   L   Y   P   D   K   G   L   K   C
76  GGC GGC CAG CGT CGC TCC ATG AAA CAA TCT GCG GTG TGT GGC TGC AAA CTT TAC CCT GAT AAG GGT CTG AAA TGT
    CCG CCG GTC GCA GCG AGG TAC TTT GTT AGA CGC CAC ACA CCG ACG TTT GAA ATG GGA CTA TTC CCA GAC TTT ACA

151 ACC AAA CGT CCG AAT CCA AAA TCG CGC GAT GAA TAC TAT GAG ATT CTG AAG TGT ATT TGC CGT GAC ACC CAG ATC
    TGG TTT GCA GGC TTA GGT TTT AGC GCG CTA CTT ATG ATA CTC TAA GAC TTC ACA TAA ACG GCA CTG TGG GTC TAG
     T   K   R   P   N   P   K   S   R   D   E   Y   Y   E   I   L   K   C   I   C   R   D   T   Q   I

226 TTC TCA AAA AGC AGT CGC AAA GAA TAT CTG AAG CGT TGC AAC AAA TTT TAC CCG TCT CTC CCG TTG TAA GCG GCC
    AAG AGT TTT TCG TCA GCG TTT CTT ATA GAC TTC GCA ACG TTG TTT AAA ATG GGC AGA GAG GGC AAC ATT CGC CGG
     F   S   K   S   S   R   K   E   Y   L   K   R   C   N   K   F   Y   P   S   L   P   L   *

301 GCA CTC GAG CAC CAC
    CGT GAG CTC GTG GTG
```

**Fig. S4.** The nucleotide and amino acid sequence of the zebrafish CXCL17 and CXCL17-like overexpressed in *E. coli*. The amino acid sequence of mature Dr-CXCL17 and Dr-CXCL17-like is shown in red, that of SmBiT in blue.

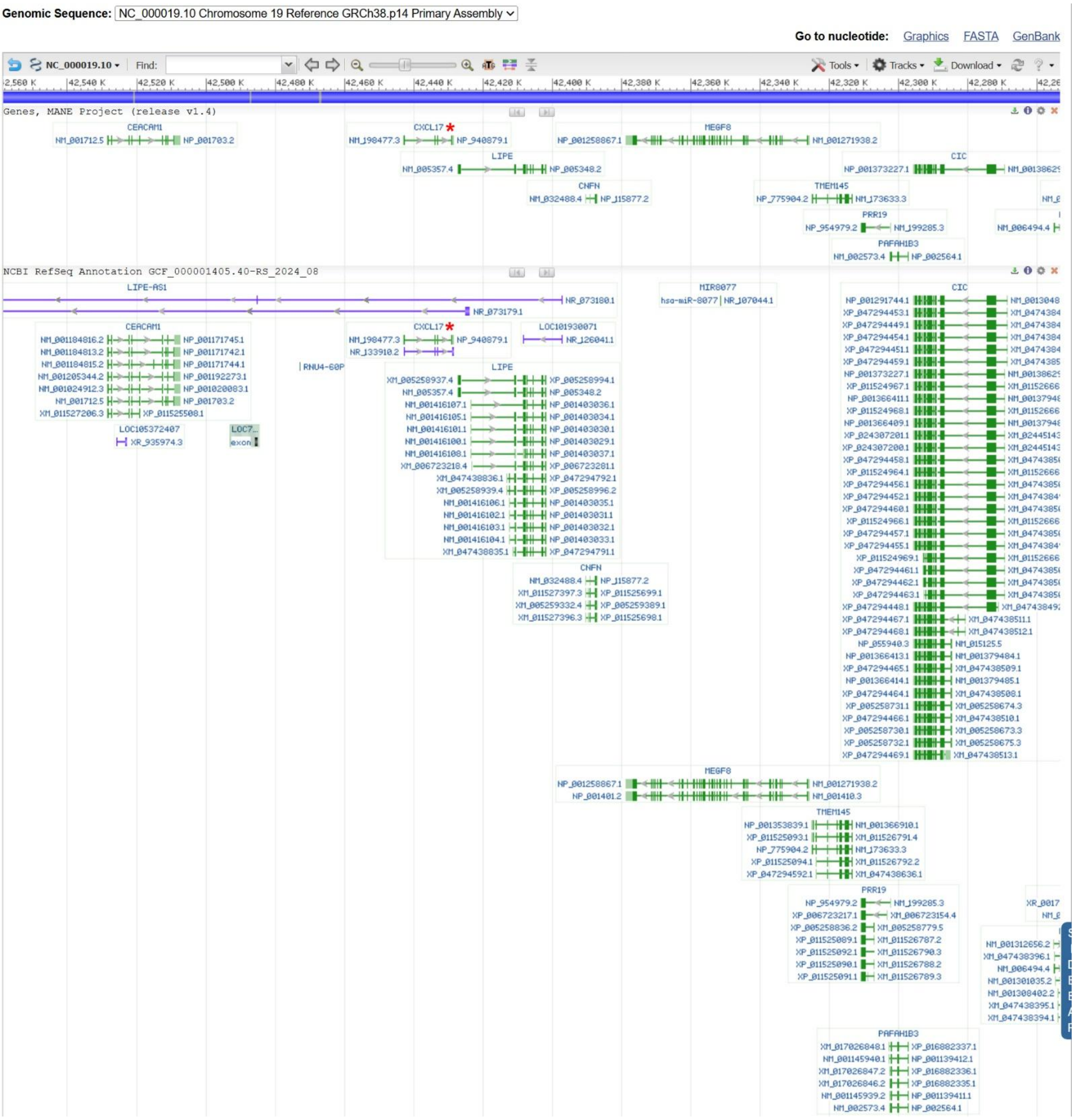


**Fig. S5.** Position of the human *CXCL17* gene in human genome. The human *CXCL17* gene is indicated by a red asterisk. The information was downloaded from the NCBI gene database (https://www.ncbi.nlm.nih.gov/gene/284340).
